# Does meiotic drive alter male mate preference?

**DOI:** 10.1101/736595

**Authors:** Sam Ronan Finnegan, Leslie Nitsche, Matteo Mondani, M. Florencia Camus, Kevin Fowler, Andrew Pomiankowski

## Abstract

Male mate preferences have been demonstrated across a range of species, including the Malaysian stalk-eyed fly, *Teleopsis dalmanni*. This species is subject to SR, an X-linked male meiotic driver, that causes the dysfunction of Y-sperm and the production all female broods. SR is associated with a low frequency inversion spanning most of the X chromosome that causes reduced organismal fitness. While there has been work considering female avoidance of meiotic drive males, the mating decisions of drive-bearing males have not been considered previously. As drive males are of lower genetic quality they may be less able to bear the cost of choice or may experience weaker selection for its maintenance if they are avoided by females. We investigated preference of drive males using binary choice trials. We confirmed that males prefer to mate with large females (indicative of greater fecundity) but found no evidence that the strength of male mate preference differs between drive and standard males. This suggests that the cost of choice does not restrict male reference among drive males. In a further experiment, we found that large eyespan males showed strong preference whereas small eyespan males showed no preference. This is likely to weaken mate preference in drive males, as on average they have reduced eyespan compared to standard males. In this respect, drive males are subject to and exert weak sexual selection.

**Lay summary:** We studied male mate preference in stalk-eyed flies. This species suffers from meiotic drive, a selfish genetic element that causes a reduction in sperm production and organismal fitness. We predicted that males with meiotic drive would show weak mate preference. Males preferred to mate with large females, but there was no difference in the strength of preference between drive and non-drive males. Males with larger eyespan showed stronger mate preference. Meiotic drive males usually have reduced eyespan so on average they exert weaker sexual selection on females, but this is mediated by eyespan, not genotype *per se*.

## Introduction

Despite a historical narrative of indiscriminate males attempting to mate with choosy females (Bateman 1948), male mate preference is established as a widespread phenomenon in sexual selection (Bonduriansky 2001; Edward and Chapman 2011). Male mate preference has even been observed in several lekking species, where males only provide sperm, across a variety of taxa including flies (Shelly et al. 2012), birds (Sæther et al. 2001) and fish (Werner and Lotem 2003). Several general conditions have been identified under which selection will favour the evolution of male mate preference (Bonduriansky 2001). The first is that mate preference must be costly, as if there are no or minimal costs, it would not pay males to be choosy (Bonduriansky 2001). Costs may arise if female sampling leads to higher predation risk, greater disease transmission or simply requires more time (Parker 1983; Pomiankowski 1987). There are also opportunistic costs since mating with one female inevitably reduces the time available to search and mate with others (Bonduriansky 2001). In addition, sperm production is costly (Dewsbury 1982) and must set limits on the mating capacity of individual males. So, males need to allocate their ejaculate strategically among females, a form of cryptic male preference (Wedell et al. 2002). On the other hand, there must be variation in female quality that affects male reproductive success, so that male choice yields a benefit (Parker 1983). The most obvious benefit that males are likely to gain from choosing among available females will arise from variation in fecundity (Bonduriansky 2001). Differences between females may be generated by current or future egg production, through variable capacity for parental investment in young, due to consequences of female age and mating status (e.g. virgin vs. mated, time since last mating, degree of sperm competition). Also females may vary in genetic quality or genetic compatibility. Overall, to promote the evolution of male mate preference the costs of assessing potential mates should be low enough that they do not outweigh the benefits of preference (Nakahashi 2008), as with female preference (Pomiankowski 1987).

The Malaysian stalk-eyed fly, *Teleopsis dalmanni*, fulfils these general conditions for the evolution of male mate preference. In the wild, male stalk-eyed flies establish lek sites at dusk on exposed root hairs, to which females are attracted, with an average of two females per lek site (range 1-7; Cotton et al. 2010). Most mating occurs in a short period (∼20-30 minutes) at dawn the next day (Burkhardt and de la Motte 1985; Chapman et al. 2005). The majority of lek aggregations contain a single male and several females (Cotton et al. 2010), providing males with the opportunity to mate selectively. The direct costs of male preference are likely to be small as females settle with the lek male, and he can easily compare them on his lek. In addition, in the dawn period there is typically no competition for mating, as only the harem male mates with females settled on his lek. However, there may be costs related to the mating rate. Mating is associated with a temporary reduction in accessory gland size, and these organs do not recover to pre-mating size for around 24 hours (Rogers et al. 2005). In an analysis of the correlates of mating frequency, Rogers et al. (2005) showed that the majority of males (76.1%) presented with six females were unable to mate with all of them within an hour (Rogers et al. 2005), and the early morning period of mating is considerably shorter than this (Cotton et al. 2010). These studies suggest that males suffer limits to their daily mating capacity, which probably extends across days. In addition, females are observed to fly off leks during the dawn period, whether they have mated or not (personal observation). A male pre-occupied mating with one female, will lose the opportunity to mate with others. Males are likely to benefit from mate preference as females vary in fecundity. In the wild, as in the laboratory, female fecundity is positively correlated with body size along with nutritional status (David et al. 1998; Cotton et al. 2010, 2015). Female eyespan is a likely target for male preference. In wild samples female eyespan is predictive of fecundity even after controlling for body size, with which it strongly covaries (Cotton et al. 2010). Indeed, male mate preference for large eyespan and high fecundity females has been reported in this species under both laboratory and field conditions (Cotton et al. 2015). Together this evidence suggests that females vary in reproductive quality in ways that will affect male fitness and the costs of male preference are unlikely to outweigh the potential benefits.

Here we go further and investigate the effect of *sex-ratio* (SR), X-linked meiotic drive, on male mate preference in *T. dalmanni*. SR systems are common in flies, where they cause male carriers to produce female-biased broods and are often associated with considerable fitness costs (Jaenike, 2001; Lindholm et al. 2016). In stalk-eyed flies, SR arises from a selfish genetic element on the X chromosome that causes dysfunction of Y-bearing sperm, and transmits itself at super-Mendelian frequencies (Johns et al. 2005). The loss of Y-bearing sperm leads to the production of strongly female-biased broods (Wilkinson and Sanchez 2001). The SR chromosome (X^SR^) exists at moderate frequencies (∼20%) in populations of *T. dalmanni* (Wilkinson et al. 2003; Cotton et al. 2014; Paczolt et al. 2017) and is estimated to have diverged from a non-driving, standard chromosome (X^ST^) approximately 500,000 years ago (Paczolt et al. 2017). The gene(s) controlling meiotic drive are located in a large paracentric inversion covering most of the X^SR^ chromosome (Johns et al. 2005; Paczolt et al. 2017), as is typical of several meiotic drive systems (Lyon 2003; Dyer et al. 2007; Pinzone and Dyer 2013). As low frequency inversions are associated with reduced recombination rates, they are subject to weaker natural selection and the accumulation of deleterious mutations (Hoffmann and Rieseberg 2008; Kirkpatrick 2010). In several drive systems, this results in reduced viability (Curtsinger and Feldman 1980; Beckenbach 1996; Larracuente and Presgraves 2012; Sutter and Lindholm 2015). There is some evidence for reduced genetic quality of SR in *T. dalmanni*. Both males and females carrying the X^SR^ chromosome have reduced egg-to-adult viability (unpublished data), even though adult longevity does not appear to differ (Wilkinson et al. 2006). In addition, SR males have repeatedly been shown to have reduced eyespan both in laboratory (Wilkinson et al. 1998, Meade et al. 2018) and wild populations (Cotton et al. 2014; Meade et al. 2018). This association probably arises from the highly condition-dependent nature of male eyespan which reflects both environmental (David et al. 1998; Cotton et al. 2004) and genetic quality (David et al. 2000; Bellamy et al. 2013).

Previous work has not investigated whether meiotic drive affects sexual preference. A number of arguments suggest that SR males will show weaker mate preference than standard (ST) males. Female mate preferences are often costly condition-dependent traits, with the highest quality females showing the strongest preference for the most attractive males (Cotton et al. 2006). For example, in the three-spined stickleback (*Gasterosteus aculeatus*), females from high condition families display strong preference for male red throat coloration while females from low condition families do not (Bakker et al. 1999). If male mate preference is costly, low condition SR males may be less able to bear this cost, leading to weaker male preferences for high value females (Howie and Pomiankowski 2018). A more direct association may arise if genes for male preference are X linked. Given greater mutational decay on the X^SR^ chromosome, SR males would be expected to display weaker preferences for the largest females. A third possibility arises from the association of SR with reduced male eyespan. Theoretical work suggests that visual perception improves as eyespan increases (Burkhardt and de la Motte 1983). Small eyespan may limit the ability of males to discriminate among female phenotypes. Such an association between eyespan and discrimination in stalk-eyed flies has been shown in female mate preference (Hingle et al. 2001), and may well extend to males. A further possibility is that since females prefer to roost and mate with males of large eyespan (Wilkinson and Reillo 1994; Wilkinson and Dodson 1997; Hingle et al. 2001; Cotton et al. 2010), then SR males on average will attract fewer females to their leks. This could result in weaker selection for mate preference among SR males if they more frequently roost in single female leks and so have less opportunity for choice. A potential example of this is the two-spotted goby, *Gobiusculus flavescens*, where large, attractive males prefer to mate with colourful females, whereas small, less attractive males express no preference despite equal courtship effort (Amundsen and Forsgren 2003).

All of these arguments lead to the prediction that SR males will show weaker mate preference than ST males. In order to test this, we used simple binary choice trials which have been used previously to measure the strength of male mate preference in stalk-eyed flies (Cotton et al. 2015). In two separate experiments, SR and ST males were presented with two females, one large and one small, and allowed to freely mate during a short time period. Two females is the mean number observed in the wild on male leks with females (Cotton et al. 2010). The design aimed to mimic, under controlled conditions, the sex ratio and time-frame under which male preference is expressed in the wild. In the first experiment, focal male eyespan was constrained to lie within a narrow range of trait values in order to test whether the genotypic differences between SR and ST males cause differences in mating behaviour independent of male eyespan. In the second experiment, focal male eyespan was unconstrained and drawn from its natural distribution so as to determine the direct effect of eyespan and its association with genotype (SR and ST) on mate preference.

## Methods

### Stocks

A stock population was obtained from Ulu Gombak in Malaysia (3°19’N 101°45’E) in 2005 (by Sam Cotton and Andrew Pomiankowski). It is maintained at 25°C on a 12:12 hour light:dark cycle at high population density. Fifteen minute artificial dawn and dusk phases are created by illumination from a single 60-W bulb at the start and end of each light phase. This population contains only standard (i.e. wildtype) males, and is referred to as the ST stock, as it does not contain individuals carrying the X^SR^ drive chromosome.

In 2012 a further collection was made of male flies from the same location (by Alison Cotton and Sam Cotton) and used to create a SR stock population that maintains the X^SR^ chromosome, following a standard protocol (Presgraves et al. 1997; Meade et al. 2018). Briefly, individual males from the SR population are housed with three ST stock females and allowed to mate freely. Offspring are collected from these crosses and the offspring sex ratio scored. Males siring female-biased broods (>90% female offspring, >15 total offspring) are considered SR (X^SR^Y), and their female progeny are therefore carriers of the SR chromosome (X^SR^X^ST^). Progeny from other males, which are likely to be ST, are discarded.

The resulting heterozygous females are then mated with ST stock males (X^ST^Y), producing SR (X^SR^Y) and ST (X^ST^Y) males in an expected 1:1 ratio. These males are crossed to three ST stock females, and the process is repeated (i.e. keeping the progeny of X^SR^Y males and discarding those of X^ST^Y males). The regular crossing with ST stock males and females homogenises the autosomes, Y chromosome, wildtype X chromosome and mitochondrial genes across the two stock populations. In other respects, the SR and ST stocks were kept under similar conditions.

### Experimental flies

Experimental males were collected from egg-lays, consisting of a petri dish containing moistened cotton wool and ∼15g pureed sweetcorn, placed into stock cages. The petri dishes were removed after 3 days and subsequently the eclosed adults were collected after 3-4 weeks. At two weeks post eclosion, flies were anaesthetised on ice and separated by sex. Experimental males (SR and ST) were collected from egg-lays taken from the SR stock population (females from those egg-plays were discarded). Experimental females were taken from the ST stock population (ST males were discarded).

Eyespan was measured as the distance between the outermost edges of the eye bulbs (Cotton et al. 2004) when flies were anaesthetised on ice using ImageJ (v1.5.0). In experiment 1, males were standardised to a narrow range of eyespan (7.5-8.5 mm) to minimise any potential effect of variation in male eyespan on female behaviour. Males were housed in large cages (35cm x 22cm x 20cm) with stock females at an even sex ratio in order for them to mate at a normal rate prior to the mating assay. Stock females used in the experiment were defined as large (eyespan ≥ 5.8 mm) or small (eyespan ≤ 5.4 mm) following Rogers et al. (2006) and Cotton et al. (2015). Intermediate size females were discarded. Large females were fed high quality food consisting of 100% pureed corn. Small females were fed low quality food consisting of 20% pureed corn and 80% sugar solution (25% sugar w/v), with the addition of an indigestible bulking agent (3% carboxymethylcellulose w/v) so the viscosity was the same as in the high quality food (Rogers et al. 2008; Cotton et al. 2015). The two diets were used to amplify differences in fecundity between the two size classes of experimental female (Cotton et al. 2015). The two classes of female were housed with stock males to allow them to mate at a normal rate.

In experiment 2, males were reared from egg-lays collected from SR stock cages with variable amounts of corn (between 1.5 – 15g). This varied the amount of food per developing larvae in order to generate size variation in eyespan and thorax. Otherwise the procedures used were the same as in experiment 1. The sole exception was that the two types of female, large and small, were fed on the same high quality food as adults.

### Male mating assays

Male flies were presented with a choice of large and small females in mating chambers (Fig. 1, Cotton et al. (2015)). Mating chambers were set up in the afternoon prior to each assay. Males were placed in the top compartment, and one large and one small female were placed in the bottom compartment. Interactions between males and females were limited during this period by a cardboard partition placed between the two compartments. At dawn on the following morning, the partition was removed, and the mating chambers were observed for 30 minutes. The number of copulations with each size class was recorded, as well as the order of mating. A successful copulation was defined as intromission lasting more than 30 seconds, as copulations shorter than this duration do not result in spermatophore transfer (Rogers et al. 2006). In some cases, the exact start or end of a copulation was not observed. If the observed duration of the mating was greater than 30 seconds, these were included in the analysis of preference. Males that attempted to mate but were unsuccessful (defined as following and attempted mounting of female, but without intromission) were presented with a different set of one large and one small female and observed for an additional 30 minutes. After completion of the assay, focal males were frozen and stored in ethanol. Females were isolated in individual 500ml pots for two days before being returned to population cages. This was to ensure that no females were used in assays on consecutive days.

**Figure 1.**
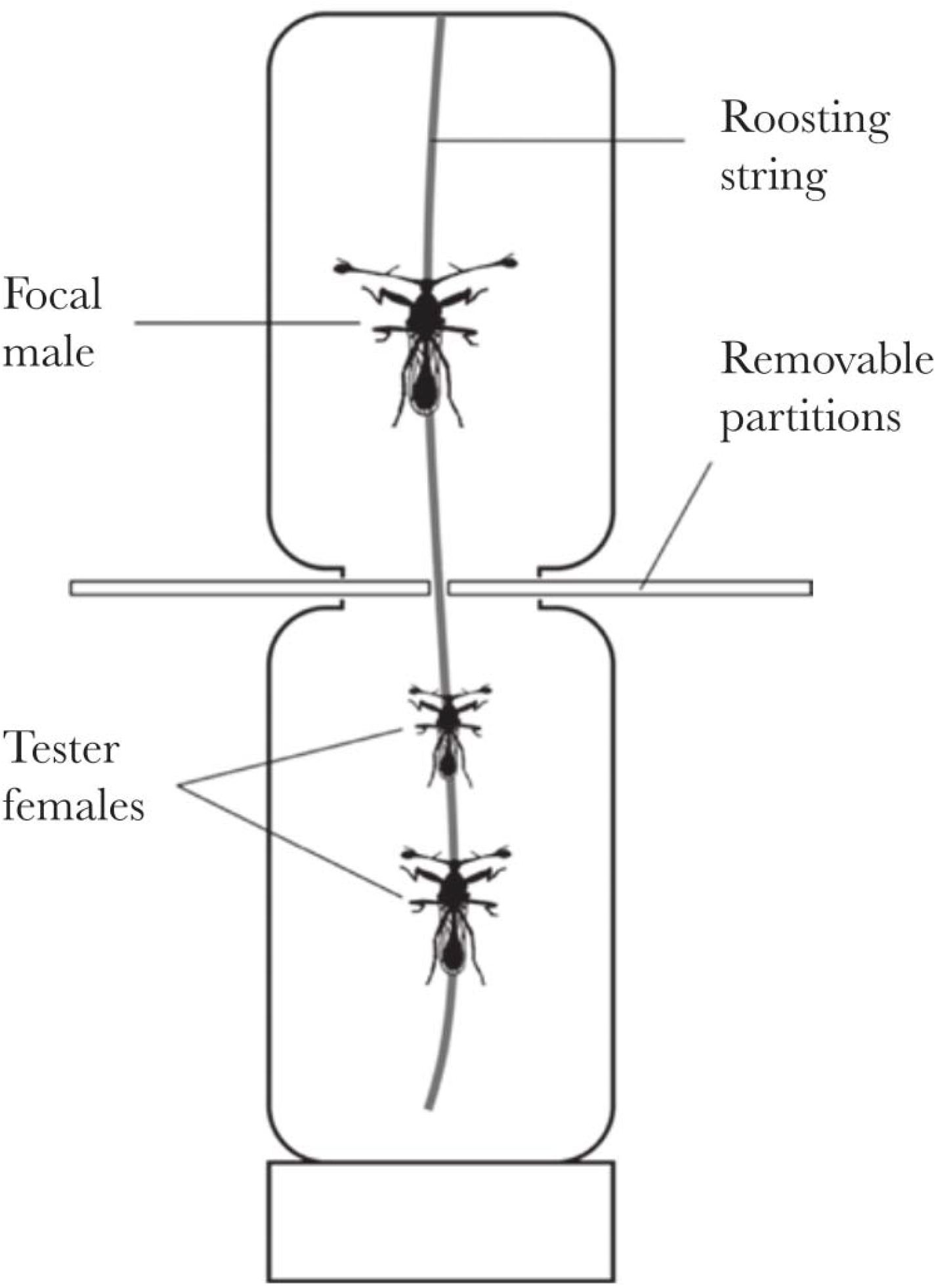
Mating chambers used for male mate preference assay. A single male of unknown genotype was placed in the top compartment, with two tester females (one large, one small) in the bottom compartment. Males and females were kept separate by a removable partition until testing commenced. A string, resembling a rootlet, runs the length of the chamber, to provide a roosting site. Reproduced with permission from Cotton et al. (2015).

### DNA extractions and genotyping

The experimenters were blind to the genotype of experimental males, as this was inferred *post-hoc* by genotyping. In experiment 1, to extract DNA, half the thorax of each fly was added to individual 1.5ml Eppendorf tubes containing 10μl Proteinase K (10mg.ml^−1^) and 250μl of DIGSOL at 55°C. Subsequently, 300μl of 4M ammonium acetate was added and samples spun down for 10 minutes at 13,000RPM. The supernatant was then aspirated into new tubes containing 1ml 100% ethanol and spun for 10 minutes at 13,000RPM to precipitate DNA. DNA was then washed in 70% ethanol to remove excess salt and left to air-dry for 45 minutes before being eluted in 30μl Low-TE (1mM Tris-HCl pH8, 0.1mM EDTA). DNA was PCR-amplified on a 2720 Thermal Cycler (Applied Biosystems) in 96-well plates with each well containing 1μl of dried DNA, 1μl of QIAGEN Multiplex PCR Mastermix (Qiagen), and 1μl of Primer mix (consisting of the forward and reverse primers for *ms395* and *comp162710* each at a concentration of 0.2μM). A drop (10μl) of mineral oil was added to limit evaporation. Fragment length analysis was carried out using an ABI3730 Genetic Analyzer (Applied Biosystems) with a ROX500 size standard. Microsatellite allele sizes were assigned using Fragman package v. 1.07 (Covarrubias-Parazan et al. 2016) in R v. 3.2.3 (R Development Core Team, 2011) and checked using GENEMAPPER 4.0. Two markers were used to distinguish SR and ST males. Microsatellite *ms395* has a bimodal distribution where large (>218bp) alleles are strongly associated with SR (Johns et al. 2005; Cotton et al. 2014; Meade et al. 2018; Paczolt et al. 2017). *Comp162710* is an indel marker with a small allele (201bp) found in SR males, and a large allele (286bp) found in ST males (Meade et al. 2018). Males with large *ms395* alleles and small *comp162710* alleles were classed as SR. Where markers gave conflicting signals, genotype was assigned on the basis of *comp162710* allele size. All PCRs and fragment length analysis were carried out at NBAF-S at the University of Sheffield.

In experiment 2, the same procedure was used to extract DNA, but a different PCR protocol was used. Each well consisted of 1μl of DNA, 0.1μl of 5x Phusion Taq polymerase (New England BioLabs), 0.2μl of dNTPs, 6.2μl UltraPure water, and 0.5μl each of the 10μM forward and reverse primers for *comp162710*. *Comp162710* fragment lengths were assayed by gel electrophoresis on a 3% agarose gel with a 0.5x TBE buffer.

### Statistical analysis – Experiment 1

We analysed the effect of genotype on the number of copulations with each size class of female using logistic regression with a quasi-binomial error structure to account for over dispersion. The intercept term in this model determines whether males show preference for either size class of females. The data was also split by genotype and the same model was run to determine if SR and ST males preferred large females. For comparison with earlier work (Cotton et al. 2015), mate preference for each individual male was assessed using an index based on the proportion of total copulations with the large female, *Pref* = (*C_L_* − *C_S_*)/(*C_L_* + *C_S_*), where *C_L_* and *C_S_* are the number of copulations with the large and small females respectively. Preference values range ±1 and are symmetric about zero. For an individual male, a value greater than zero indicates preference for large females, and less than zero indicates preference for small females. Preference in each consecutive mating was assessed using binomial tests on the number of copulations with large and small females, on the pooled dataset, and SR and ST males separately. The effect of genotype on the number of copulations with large and small females was analysed on each consecutive mating using generalised linear models with quasibinomial error distributions.

### Statistical analysis – Experiment 2

Experiment 2 was designed to determine whether male eyespan had an effect on mating preference, and whether this relationship varied according to male genotype. First, the effect of male eyespan, genotype and their interaction were modelled on the number of copulations with each size class of female in a generalized linear model with a quasi-binomial error structure. Then, males were split into three groups on the basis of their eyespan: small (eyespan < 6.0mm), medium (eyespan 6.0mm - 7.5mm) and large (eyespan > 7.5mm). The effect of eyespan group, genotype, and their interaction on the number of copulations with each size class of female was analysed in a generalised linear model with a quasi-binomial error distribution. Difference in mean preferences of each size group was assessed using the glht function of the multcomp package in R. The effect of genotype on thorax length and eyespan was analysed in a linear model. Preference in each consecutive mating was analysed as above. The effect of eyespan and genotype on the probability of mating at all was analysed in a binomial generalised linear model, as above.

## Results

### Experiment 1 - Genotype and male preference

In the first experiment, males showed a preference for large females when genotypes were pooled (*Pref* mean ± SE = 0.3721 ± 0.055; *t* = 6.413, *P* < 0.0001, *n* = 162). Males preferred large females in their first (*Pref* mean ± SE = 0.4321 ± 0.0711, *P* < 0.0001, *n* = 162), second (*Pref* mean ± SE = 0.3030 ± 0.0832, *P* = 0.0006, *n* = 132) and third mating (*Pref* mean ± SE = 0.4257 ± 0.0904, *P* < 0.0001, *n* = 101). For subsequent matings there was no male preference for large females, in large part reflecting the reduced sample size (fourth mating: *Pref* mean ± SE = 0.1803 ± 0.1269, *n* = 61, *P* = 0.2000; fifth mating: *Pref* mean ± SE = 0.2593 ± 0.1894, *n* = 27, *P* = 0.2478).

The preference of SR and ST males did not differ from each other (GLM: *t* = −0.353, *P* = 0.725, *n* = 157). Preference was for large eyespan females in both SR (*Pref* mean ± SE = 0.4145 ± 0.079, *t* = 4.848 *P* < 0.0001, *n* = 81) and ST males (*Pref* mean ± SE = 0.3367 ± 0.0806, *t* = 4.053, *P* = 0.0001, *n* = 76; Figure 2). Across consecutive copulations, SR and ST males preferred large females in the first (SR *Pref* mean ± SE = 0.5062 ± 0.0964, *P* < 0.0001, *n* = 81; ST *Pref* mean ± SE = 0.3684 ± 0.1073, *P* = 0.0018, *n* = 76), second (SR *Pref* mean ± SE = 0.3333 ± 0.1227, *P* = 0.013, *n* = 60; ST *Pref* mean ± SE = 0.2647 ± 0.1178, *P* = 0.0385, *n* = 68), and third (SR *Pref* mean ± SE = 0.3000 ± 0.1526, *P* = 0.0807, *n* = 40; ST *Pref* mean ± SE = 0.4737 ± 0.1177, *P* = 0.0005, *n* = 57) mating, and did not differ in the strength of their preference across these copulations (1^st^ mating *F*_1,155_ = 0.9107, *P* = 0.3414; 2^nd^ mating *F*_1,126_ = 0.1623, *P* = 0.6878; 3^rd^ mating *F*_1,95_ = 0.8226, *P* = 0.3667).

**Figure 2.**
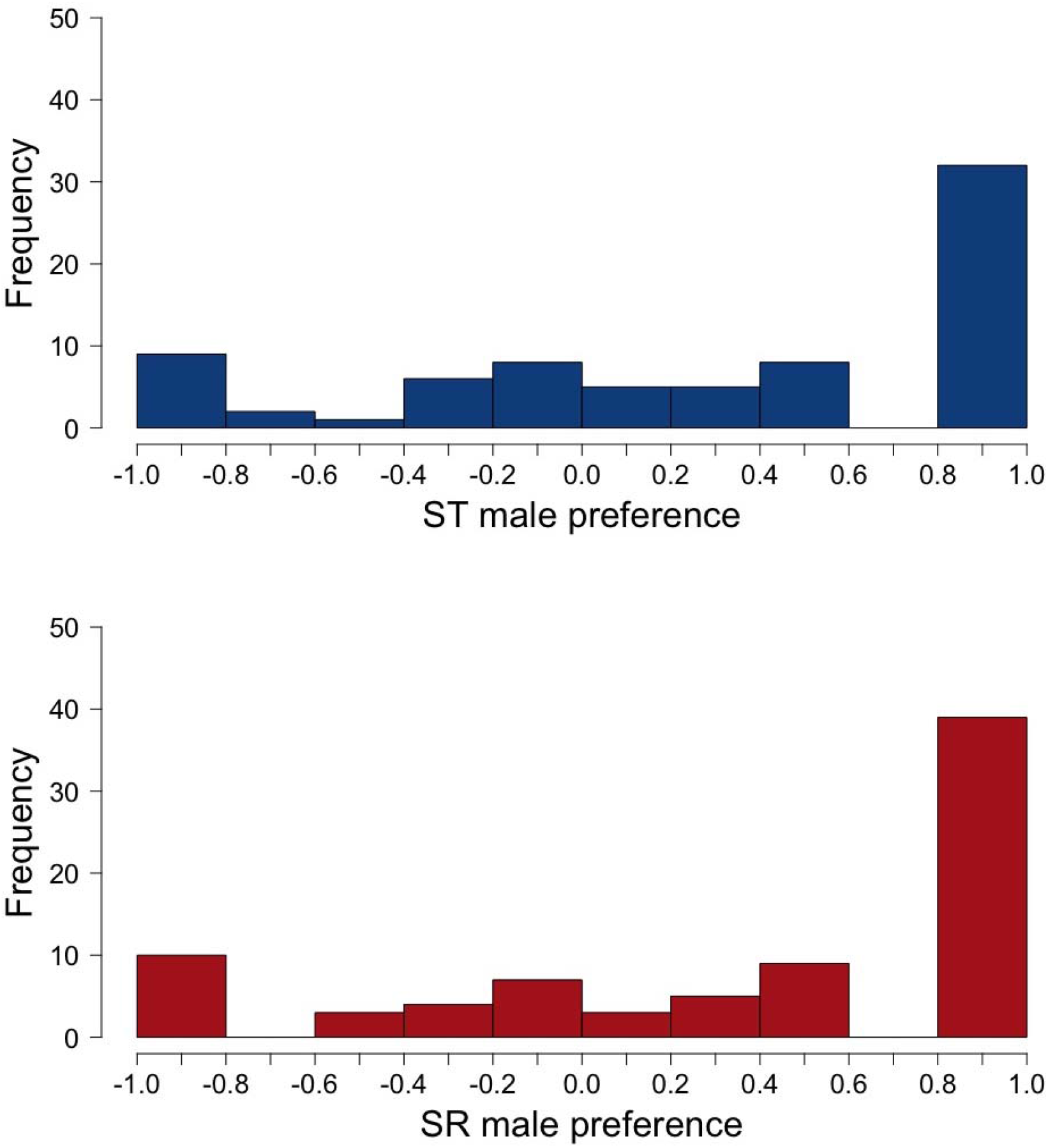
Frequency distribution of mate preference values for ST (top) and SR (bottom) males from experiment 1. Bins are right-closed, left-open. So for example, males with zero preference fall within the bin −0.2 < *x* ≤ 0.0).

### Experiment 2 - Eyespan and male preference

In the second experiment, larvae were exposed to variable amounts of food during development. Adult males showed considerable variation in eyespan (mean ± SD = 7.026 ± 1.495 mm, range 3.625 – 9.461 mm). Eyespan was strongly co-linear with body size (i.e. thorax length, *F*_1,191_ = 788.5, *P* < 0.0001), but did not differ with genotype (*F*_1,191_ = 0.9322, *P* = 0.3355), nor was there a difference in the allometric slope of eyespan on body size with genotype (*F*_1,191_ = 0.0014, *P* = 0.9706).

As before, when individuals from both genotypes were pooled, males showed a preference for large females overall (*Pref* mean ± SE = 0.2344 ± 0.0494, GLM: *t* = 5.922, *P* < 0.0001, *n* = 178), and in the first (*Pref* mean ± SE = 0.3371 ± 0.0707, *P* < 0.0001, *n* = 178), second (*Pref* mean ± SE = 0.2785 ± 0.0767, *P* = 0.0005, *n* = 158) and third matings (*Pref* mean ± SE = 0.2500 ± 0.083, *P* = 0.0033, *n* = 135). Again, there was no male preference for large females in subsequent matings as sample size fell (fourth mating, *Pref* mean ± SE = 0.1132 ± 0.0970, *P* = 0.2853, *n* = 107; fifth mating, *Pref* mean ± SE = 0.2500 ± 0.1220, *P* = 0.0599, *n* = 64).

Male eyespan had a strong positive effect on mating preference (*F*_1,174_ = 9.2135, *P* = 0.0027, Figure 3). When males were split into three groups based on eyespan (large >7.5mm, medium 6.0 – 7.5mm and small <6.0mm), male eyespan group (*F*_2,173_ = 3.5760, *P* = 0.0009, *n* = 197) affected preference, with larger males showing stronger preference than medium (|Z| = 2.754, *P* = 0.0158) and small males (|Z| = 3.430, *P* = 0.0018). Large males preferred to mate with large females (*Pref* mean ± SE = 0.4110 ± 0.0309, *t* = 3.840, *P* = 0.0003, *n* = 89). Medium males (*Pref* mean ± SE = 0.1919 ± 0.0828, *t* = 1.910, *P* = 0.0611, *n* = 63) and small males showed no preference (*Pref* mean ± SE = −0.0702 ± 0.1263, *t* = 0.4040, *P* = 0.6880, *n* = 50).

**Figure 3.**
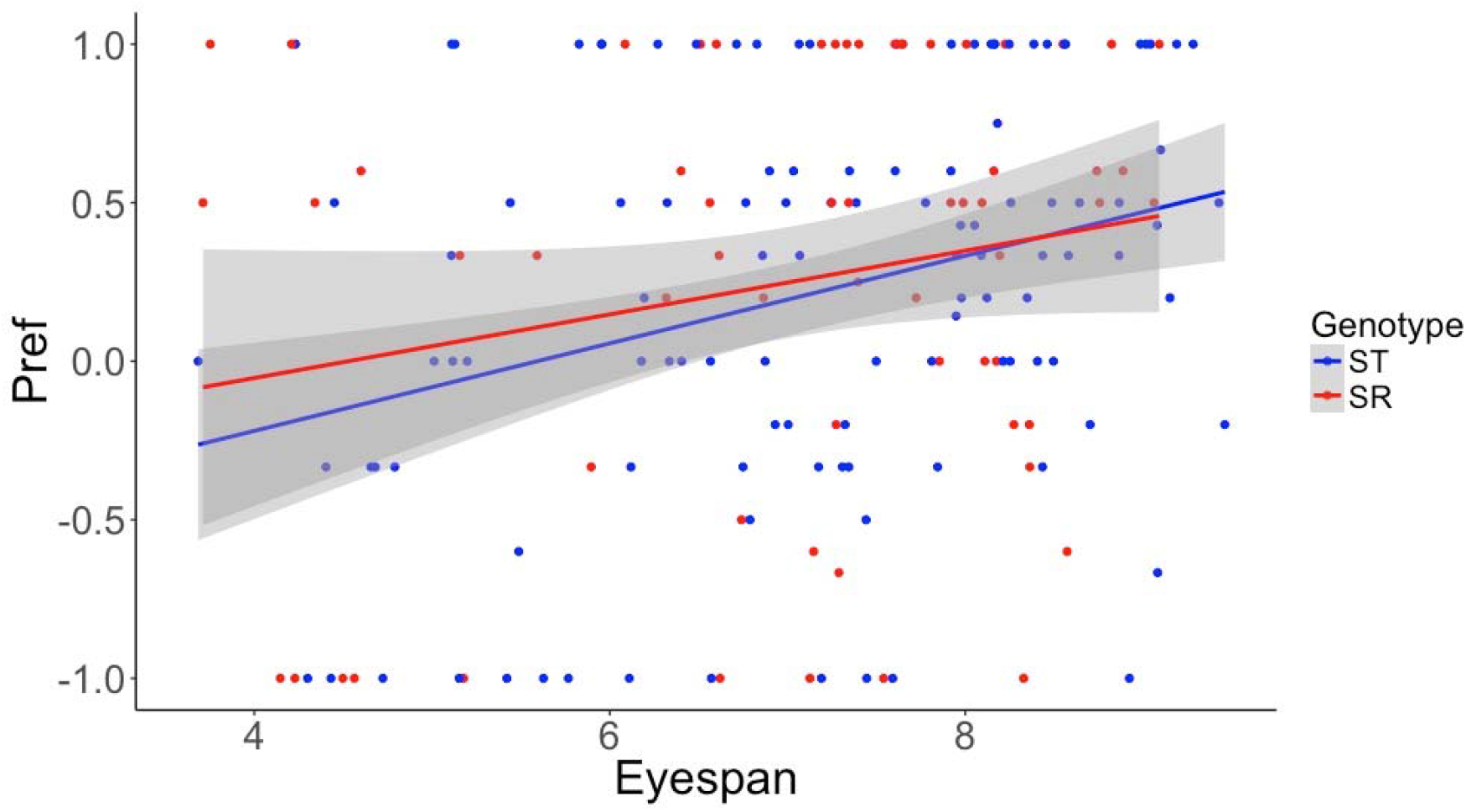
Line graph showing a regression of *Pref* values on eyespan for ST and SR males. Shaded areas represent 95% confidence intervals.

As in the first experiment, there was no difference in the strength of preference according to genotype (*F*_1,171_ = 0.6282, *P* = 0.4291). Both SR (*Pref* mean ± SE = 0.2508 ± 0.0887, *t* = 3.621, *P* = 0.0006, *n* = 69) and ST males (*Pref* mean ± SE = 0.2156 ± 0.0600, *t* = 4.450, *P* < 0.0001, *n* = 128) preferred large females. After controlling for the effect of eyespan group (large, medium, small eyespan), there was still no effect of genotype on the strength of preferences (all *P* > 0.4), nor any interaction between eyespan group and genotype (*F*_2,170_ = 0.1522, *P* = 0.8589). Both SR and ST males preferred large females in the first (SR *Pref* mean ± SE = 0.3871 ± 0.1181, *P* = 0.0044, *n* = 61; ST *Pref* mean ± SE = 0.2982 ± 0.0898, *P* = 0.0019, *n* = 114), second (SR *Pref* mean ± SE = 0.4286 ± 0.1218, *P* = 0.0018, *n* = 56; ST *Pref* mean ± SE = 0.1800 ± 0.0988, *P* = 0.0066, *n* = 100), and third (SR *Pref* mean ± SE = 0.3191 ± SE 0.1397, *P* = 0.0011 *n* = 47; ST *Pref* mean ± SE = 0.2093 ± 0.1061, *P* = 0.0007, *n* = 86) mating, and there was no difference in the strength of SR and ST preference across these matings (1^st^ mating *F*_1,174_ = 0.3541, *P* = 0.5525; 2^nd^ mating *F*_1,154_ = 2.4044, *P* = 0.1230; 3^rd^ mating *F*_1,131_ = 0.3874, *P* = 0.5437).

## Discussion

Male mate preferences have been observed across a range of species, even where initially unexpected, for example in polygynous species which lack paternal care or other forms of direct male investment in offspring or mating partners (Edward and Chapman 2011). In this study of stalk-eyed flies, we found that males show preference for large eyespan females. This mirrors previous laboratory and field studies in *T. dalmanni* (Cotton et al. 2015). As in other species, the likely benefit of this preference lies in mating with higher fecundity females (Olsson 1993; Dosen and Montgomerie 2004; Byrne and Rice 2006; Reading and Backwell 2007). 2015). Female eyespan is a reliable indicator of fecundity among field caught stalk-eyed flies, where it explains a significant amount of variation in ovarian egg number, even after controlling for body size (Cotton et al. 2010, 2015).

There was no difference between SR and ST males in their strength of preference. In order to compare genotypes independent of differences in size, in the first experiment male eyespan was restricted to a narrow range at the large end (7.5-8.5mm) of the distribution. Male eyespan is a highly condition-dependent trait, sensitive to both environmental and genetic differences (David et al. 2000, Cotton et al. 2004, Bellamy et al. 2013). By placing limits on the eyespan of experimental males, we may have inadvertently picked out SR and ST males of equivalent high condition and masked differences between the genotypes. This may be a problem as SR is associated with a large chromosomal inversion covering most of the X (Johns et al. 2005), and is predicted to have accumulated deleterious alleles (Hoffmann and Rieseberg 2008; Kirkpatrick 2010) and is known to be associated with reduced eyespan (Cotton et al. 2014; Meade et al. 2018). So, concentrating on large flies may even have selected SR males with higher condition than ST males. To address this concern, a second experiment collected males that eclosed from eggs laid on variable quantities of food. This procedure resulted in a much greater range in male eyespan among experimental males, with both smaller and larger eyespan (3.6-9.5mm). Again, as with the first experiment, there was no difference in the strength of mate preference between SR and ST males. Nor were there any preference differences between SR and ST males that had small, medium or large eyespan.

The two experiments are similar but not clones of each other. As well as the differences already mentioned in the eyespan range of experimental male, there was a minor in the tester females. In the first experiment, small females were fed a low value diet known to decrease egg production, and large females were fed a high value diet known to increase egg production (Cotton et al. 2015). In the second experiment, large and small eyespan females were fed the same diet, reducing fecundity difference. Previous work shows that males independently prefer females with large eyespan and females with high fecundity (Cotton et al. 2015). There was still male preference for the large eyespan females and no difference in preference between SR and ST males in the second experiment despite the lack of dietary manipulation. Having two experiments with slightly different protocols yielding the same result lends greater credence to our claim that male preference does not differ between SR and ST males.

We deliberately designed the experiment to simulate the field behaviour of stalk-eyed flies. In the wild, leks form at dusk, attract a restricted number of females (mean 2, range 1-7) and are the sites where most copulations take place at dawn the following day (Cotton et al. 2010). The experimental protocol tracked males for 30 minutes at dawn, allowing males to mate multiply and exert mate preference between a large and small female. Our design presented males with a binary choice of large and small females and this is appropriate given the biology of stalk-eyed flies. Preference assessments based on choices made between two markedly different phenotypes have been criticised for a number of reasons, in particular that this approach fails to capture a “preference function” based on response to the full range of female phenotypes (Wagner 1998; Cotton et al. 2006). However, there is no particular reason to believe this would impact preferences differently in SR and ST males. In one respect, our design is unrepresentative of natural behaviour, as females leave lek sites once they have mated and females do not mate multiple times with the same male (Cotton et al. 2015). The mating chamber’s design precluded female departure. But the lack of female departure and female re-mating do not appear to prejudice the findings as, in both experiments, there was no difference between SR and ST male preference in favour of the large females in the first, second and third matings. It seems unlikely that our design masked differences in male mate preference between the two genotypes.

While there was no difference in the preference of SR and ST males, we did find that eyespan affected male preference for large females, with large eyespan males showing the strongest preference and small eyespan males exhibiting no preference at all. Vision is the dominant sensory mode for assessment of potential mates in stalk-eyed flies (Chapman et al. 2005; Chapman et al. 2017), and both stereoscopic vision and visual acuity are improved as eyespan increases (Burkhardt and de la Motte, 1983; de la Motte and Burkhardt, 1983). It follows that males with larger eyespan will be better able to distinguish differences between females and express stronger preference. Just such a relationship has been reported for female mate preference in *T. dalmanni* (Hingle et al. 2001). As mean male eyespan is smaller in SR than ST males (Wilkinson et al. 1998; Cotton et al. 2014; Meade et al. 2018), SR males will exert weaker preference. In addition, field samples show that males with smaller eyespan attract fewer females to their lek sites (Cotton et al. 2010). So, on average SR males will less frequently attract multiple females to their leks, and so have fewer opportunities for choice. The extent of this disadvantage needs to be assessed with field data to establish how lek size varies with male genotype. Finally, we note that in the second experiment, males were occasionally observed attempting to mount large females, and these females moved away or actively rejected the mating attempt by displacing their genitalia or pushing the male away. These observations are too few for statistical analysis, but it is of interest that most instances (8/10) of female rejection behaviour were directed towards small males, the remainder towards medium sized males (2/10) and none to large males.

Originally we predicted that SR males would have weak preference if male choice is costly and condition-dependent, but this is not supported by the experimental data. This weak SR preference hypothesis is based on the expected accumulation of deleterious alleles on the X^SR^chromosome inversion leading to SR males having lower genetic quality than ST males. It seems unlikely that there is no mutation accumulation in the X^SR^ inversion(s) in *T. dalmanni*, although the extent to which it impacts larval and adult fitness needs to be established in greater depth across a range of environmental conditions. A more plausible explanation is that the expression of male preference is neither costly nor condition-dependent. The absence of male-male competition at dawn when most mating takes place (Cotton et al. 2010) and the short amount of time before female lek departure do not point to obvious male preference costs associated with distinguishing between females that have already settled at a lek site. SR males may have fewer opportunities to choose between females as they fail to develop the attractive phenotype (large eyespan) used by females in deciding which lek site to join. Smaller eyespan may also mean that SR males lose out to rival males in establishing ownership of favourable lek sites and keeping other males from interloping and sneaking copulations during the dusk aggregation period. But when more than one female settles, SR males likely benefit from preferential mating with large females (leading to fecundity benefits), just as ST males do.

This is the first attempt to measure how meiotic drive influences male mating preference. We showed that male stalk-eyed flies, a harem-based polygynous species, show consistent preferences for large females, closely replicating earlier findings. There was no weakening in the strength of male preference caused by the predicted reduction in genetic condition associated with meiotic drive. There was, however, a reduction in preference as eyespan decreased, likely due to a reduction in the ability to differentiate between females of different size. This is likely to affect drive males more, as they on average have reduced male eyespan. To fully gauge the impact of these findings, further work will endeavour to understand how the expected reduced attractiveness of drive males impacts their ability to attract females and so exercise mate choice.

